# The association of extracellular vesicle (EV)-cargo miR-330-3p with postoperative delirium and a potential mechanism of tau phosphorylation and neuron toxicity

**DOI:** 10.64898/2026.03.30.715460

**Authors:** Tomonari Fujimori, Sushmita Chakraborty, Atsushi Miyagawa, Harshita Tak, Atsushi Yamaguchi, Charles W. Hogue, Charles H. Brown, Samarjit Das

## Abstract

**Background:** Postoperative delirium (POD) is a frequent and severe neurocognitive complication following cardiac surgery, associated with poor long-term outcomes. The underlying mechanisms are unclear, and objective biomarkers are urgently needed.

**Methods:** We used pre- and post-operative plasma samples from 59 patients undergoing cardiac surgery in three separate studies with rigorous delirium assessment using the Confusion Assessment Method in a case-control design. Small extracellular vesicles (sEVs) were isolated from plasma, and their miRNA cargo was profiled using RNA sequencing. Target miRNAs were validated by qRT-PCR, and digital PCR (dPCR). The functional impact of the lead candidate miRNA was investigated *in vitro* by assessing tau phosphorylation and cell viability in HT22 neuronal cell line.

**Results:** There were no differences in sEV morphology or numbers between patients with and without POD. While three candidate miRNAs were initially validated by qRT-PCR, subsequent dPCR analysis confirmed that only the perioperative change in plasma sEV-cargo miR-330-3p expression was significantly greater in patients who developed POD (*n =* 20) compared with those who did not (*n =* 20) (5.22 copies/μL plasma; 95% Confidence Interval (CI), 1.187 to 9.256; *p* = 0.0139). Receiver operating characteristic curve analysis for this change yielded an area under the curve of 0.745 (95% CI, 0.589 to 0.901). *In vitro* overexpression of miR-330-3p in a neuronal cell line significantly increased the phosphorylation of tau at Ser199 (*p* < 0.0001) and Ser396 (*p* < 0.001) and reduced cell viability (*p* < 0.001).

**Conclusions:** Our findings suggest that sEV-bound miR-330-3p increases in patients with POD after cardiac surgery. *In vitro* results suggest a potential pathogenic role for miR-330-3p, linking a systemic signal to tau-related neuronal injury.

**Clinical Perspective:** *What Is New?:*

- This study identifies a specific perioperative increase in small extracellular vesicle (sEV)-cargo miR-330-3p in patients with postoperative delirium (POD) following cardiac surgery.
- We provide the first evidence that miR-330-3p directly induces tau hyperphosphorylation and reduces neuronal viability *in vitro*, establishing a potential mechanistic link between systemic sEV signaling and neurodegeneration.

*What Are the Clinical Implications?:*

- The measurement of perioperative change in miR-330-3p could serve as an objective biological marker to assist in the early identification and risk stratification of patients at high risk for POD.
- The identified miR-330-3p/tau pathway represents a potential new therapeutic target; future interventions aimed at inhibiting this specific miRNA might help prevent or mitigate POD-related neuronal injury.
- These findings emphasize the importance of monitoring dynamic sEV-cargo changes to better understand and manage perioperative neurocognitive disorders.

## INTRODUCTION

Coronary artery bypass grafting (CABG) and other cardiac surgical procedures are among the most common major operations performed globally in older adults. A frequent complication of cardiac surgery is postoperative delirium (POD), a form of acute brain dysfunction characterized by a disturbance in attention, awareness, and cognition.^1^ The incidence of delirium after cardiac surgery is reported to be between 16-56% ^2, 3^, but is frequently underdiagnosed in clinical practice, particularly when it presents as the hypoactive subtype.^2, 3^ Delirium has been independently associated with increased long-term mortality ^4^, prolonged duration in the intensive care unit (ICU) and hospital, higher hospital charges ^3^, and a higher likelihood of both functional ^5^ and long-term cognitive decline after discharge.^6, 7^

The pathophysiology of POD is likely multifactorial, resulting from a complex interaction of patient-specific vulnerabilities (e.g., advanced age, preexisting cognitive impairment) and perioperative insults.^8^ While the precise mechanisms remain to be fully elucidated, a leading hypothesis suggests that the systemic inflammatory response to surgery can lead to neuroinflammation and disruption of the blood-brain barrier.^1, 9^ This inflammatory response, thought to involve pro-inflammatory cytokines, can trigger or accelerate neuropathological processes. However, the specific molecular signals that connect systemic insults to central nervous system dysfunction after surgery remain poorly defined.^1^ Further, the lack of objective biological markers of delirium leads to a reliance on subjective bedside assessments, which has hampered efforts to predict, prevent, and treat POD effectively.^10^

Small extracellular vesicles (sEVs) are any functional cell-secreted, membrane-bound particles that mediate interorgan communication, potentially through cargo that include microRNAs (miRNAs).^11, 12^ miRNAs are small non-coding RNA molecules that are powerful post-transcriptional regulators of gene expression.^13^ Circulating miRNAs in the blood have been shown to mediate communication between distant cells and have been implicated in the pathophysiology of several disease states.^12, 14, 15^ Although sEVs and their miRNA cargo can cross the blood-brain barrier, a potential role for circulating miRNA in the pathogenesis of POD has not been established.

In this study, we examined a role for sEV-cargo miRNA signaling in the pathophysiology of delirium after cardiac surgery. We hypothesized that the systemic insult of cardiac surgery would alter the miRNA profile differentially in cardiac surgery patients who developed versus those who do not develop POD. Using the results of this profiling, we further examined the functional impact of one differentially expressed miRNA on tau phosphorylation and cell viability in a neuronal cell model.

## Material and Methods

### Ethical Approval

This study was approved by the Johns Hopkins Hospital Institutional Review Board (Baltimore, MD, USA; IRB00030360, IRB_NA00027003, IRB00086547), and written informed consent was obtained from all participants. The study was conducted in accordance with the principles of the Declaration of Helsinki.

### Study Design, Participants, and Delirium Assessment

To control for potential confounding factors, patients who developed POD cases were matched with patients who did not develop POD (controls) from several patient cohorts, using a multi-stage experimental design. First, an initial, unbiased screening by RNA-sequencing was performed on available samples with and without POD from a discovery cohort (*n =* 6 per group), derived from the control arm of a pilot trial to examine the effects of remote ischemic preconditioning on POD or cardiac surgery induced acute kidney injury (AKI).^2, 16^ Eligibility criteria for patients in this study included age>65 years old and undergoing coronary artery bypass and/or valve cardiac surgery. Based on the results of the initial screening, candidate miRNAs were validated by qRT-PCR using a combined cohort (*n =* 10 non-POD, *n =* 8 POD). These samples were derived from the initial discovery cohort and additional available samples from a separate trial of cerebral autoregulation monitoring, with eligibility criteria that included age>55 years old at high risk for neurological complications ^17, 18^. Patients from this latter trial were matched based on age +/-5 years and were all male and part of the control arm of the study. Finally, dPCR (*n =* 20 per group) was performed as a larger, independent validation to ensure robustness of the findings. Available samples for the dPCR validation were derived from an ongoing cohort study with eligibility criteria that included age>55 years old and undergoing coronary artery bypass surgery. Patients from this cohort were matched based on sex and age +/-5 years. To confirm that any changes in miRNA expression between POD groups was not due to increased size or number of exosomes, we also used a subset of samples from this latter cohort for sEV characterization. For all cohorts, POD was assessed using the Confusion Assessment Method by experienced research staff.

### Blood Sample Collection and Plasma Processing

Whole blood was collected into ethylenediaminetetraacetic acid (EDTA) tubes at two time points: baseline and on the first postoperative day. Plasma was separated by centrifugation, and the resulting cell-free plasma was aliquoted and stored at −80°C until analysis.

### Analysis of Small Extracellular Vesicles (sEVs) from Patient Plasma

#### Exosomes or sEVs Isolation

sEV fraction was isolated from 200 μL of plasma using differential centrifugation, including ultra centrifugation method.

#### sEVs Characterization

To characterize the sEVs extracted from plasma, we utilized the Leprechaun platform (Unchained Labs, Pleasanton, CA, USA). sEVs were incubated on a microarray of antibody spots targeting CD63, CD81, CD9, or mouse immunoglobulin G (MIgG, a non-specific binding control). The concentration, size, and tetraspanin profile of isolated sEVs were characterized using Single Particle Interferometric Reflectance Imaging Sensor (SP-IRIS) with multiplex immunofluorescence for CD9, CD63, and CD81, according to the manufacturer’s protocol.

#### RNA Isolation and Sequencing

miRNA-enriched RNA was isolated from sEVs using the miRNeasy Serum/Plasma Advanced Kit (Cat#217204, Qiagen, Valencia, CA, USA). RNA libraries were prepared using the QIAseq miRNA Library kit (Cat#331505, Qiagen, Valencia, CA, USA) and sequenced on a NextSeq platform (Illumina, San Diego, CA, USA). Primary data analysis was performed using GeneGlobe software (Qiagen, Valencia, CA, USA).

#### Quantitative Real-Time PCR (qRT-PCR)

Reverse-transcribed RNA was processed using miScript Reverse Transcription Kit (Cat#218161, Qiagen, Valencia, CA, USA). PCR was performed using miScript SYBR Green Kit (Cat#218073, Qiagen, Valencia, CA, USA) at QuantStudio 5 Real-Time PCR (Thermo Fisher Scientific). All reactions were carried out in triplicate. The miRNA primers, miR-1307-5p, miR-let-7f-2-3p, miR-1245a, miR-127-5p, miR-1271, miR-136-5p, miR-17-3p, miR-222-5p, miR-330-3p, miR-671-3p, and SNO RD61 were purchased as miSript Primer Assay (Cat#218300, Qiagen, Valencia, CA, USA).

#### Digital PCR (dPCR)

For absolute quantification of validated miRNAs, cDNA was synthesized from sEV-derived RNA using the miRCURY LNA RT Kit (Cat#339340, Qiagen, Valencia, CA, USA). The copy number of each miRNA (let-7f-2-3p, miR-136-5p, miR-330-3p) was determined using the QIAcuity One dPCR System (Qiagen, Valencia, CA, USA). The assays used were the miRCURY LNA miRNA PCR Assays (Cat#339306, Qiagen, Valencia, CA, USA). Final concentrations were calculated as copies/μL of plasma (**Supplementary Information 1**).

### *In Vitro* Functional Studies

#### Cell Culture and miR-330-3p overexpression

The HT-22 mouse hippocampal neuronal cell line (Sigma-Aldrich, St. Louis, MO) was used for functional experiments. Cells were transfected for 48 hours using Lipofectamine RNAiMAX Reagent (Cat#13778500, Thermo Fisher Scientific, Waltham, MA, USA) with either scramble (Scr) control or hsa-miR-330-3p mimic (both supplied as miRCURY LNA miRNA Mimics, Cat#339173, Qiagen, Valencia, CA, USA).

#### RNA Isolation and qRT-PCR from cell lysate

Successful transfection was confirmed by isolating total RNA using the miRNeasy Mini Kit (Cat#217004, Qiagen, Valencia, CA, USA) followed by qRT-PCR quantification of miR-330-3p expression. SNO-RD61 (Cat#218300, Qiagen, Valencia, CA, USA) was used to normalize miRNA expressions from the HT-22 lysates.

#### Western Blot Analysis

The effects of miR-330-3p overexpression on tau phosphorylation were assessed by Western blot using primary antibodies against phospho-Tau (Ser199), phospho-Tau (Ser396), total Tau, and GAPDH. Densitometric analysis was performed with ImageJ.

#### Cell Viability Measurement

To identify the effect of miRNA-330-3p overexpression on HT-22 viability, the colorimetric dimethylthiazol (MTT) assay was performed. The CyQUANT MTT Cell Viability Assay Kit (Cat#V13154, Thermo Fisher Scientific, Waltham, MA, USA) was used following to the company’s instructions. Briefly, after 48 hours of transfection with either Scr or miR-330-3p mimic, MTT dye was added into the cell culture for 3 hours. The formazan crystals were then solubilized with DMSO and the absorbance was measured using a FLUOstar Omega (BMG LABTECH, Cary, NC) at 570 nm.

#### Statistical Analysis

Data distribution was assessed for normality using the Shapiro-Wilk test. For the comparison of baseline patient characteristics, continuous variables were compared using the Mann-Whitney U test, and categorical variables were compared using Fisher’s exact test. For the primary analysis of RNA-sequencing data, differential expression was calculated using GeneGlobe software (Qiagen). miRNAs were considered candidates for subsequent validation if they demonstrated a fold change of >2 (upregulation) or <0.5 (downregulation). Student’s *t*-test was used for qRT-PCR analysis. For this analysis, we excluded values (<5% in total) that did not meet technical specifications (i.e. undetectable count [*n* = 4 samples] or extreme outliers with concentrations >10-fold that of the group mean [*n* = 3 samples]). Similarly, we excluded 1 patient in the comparison of sEV particle concentration because it did not meet technical specifications. For the dPCR data with two timepoints, a two-way repeated measures ANOVA followed by Sidak’s multiple comparisons test was used. The comparison of the perioperative change in miR-330-3p concentration between matched pairs was performed using a paired *t*-test. For the *in vitro* experiments with three groups, a one-way ANOVA followed by Tukey’s post-hoc test was used. Receiver operating characteristic (ROC) curve analysis was used to evaluate the predictive utility of the perioperative change in miR-330-3p concentration for POD. A *P*-value < 0.05 was considered statistically significant. All analyses were performed using Prism 8 (GraphPad Software) or EZR.^19^

Detailed methods and data are provided in the *Online Supplemental*.

## Result

The overall workflow of the study is depicted in **Figure 1**. Our primary analysis focused on comparing sEV-miRNA profiles between the two clinical outcome groups (POD and non-POD).

**Figure 1.**
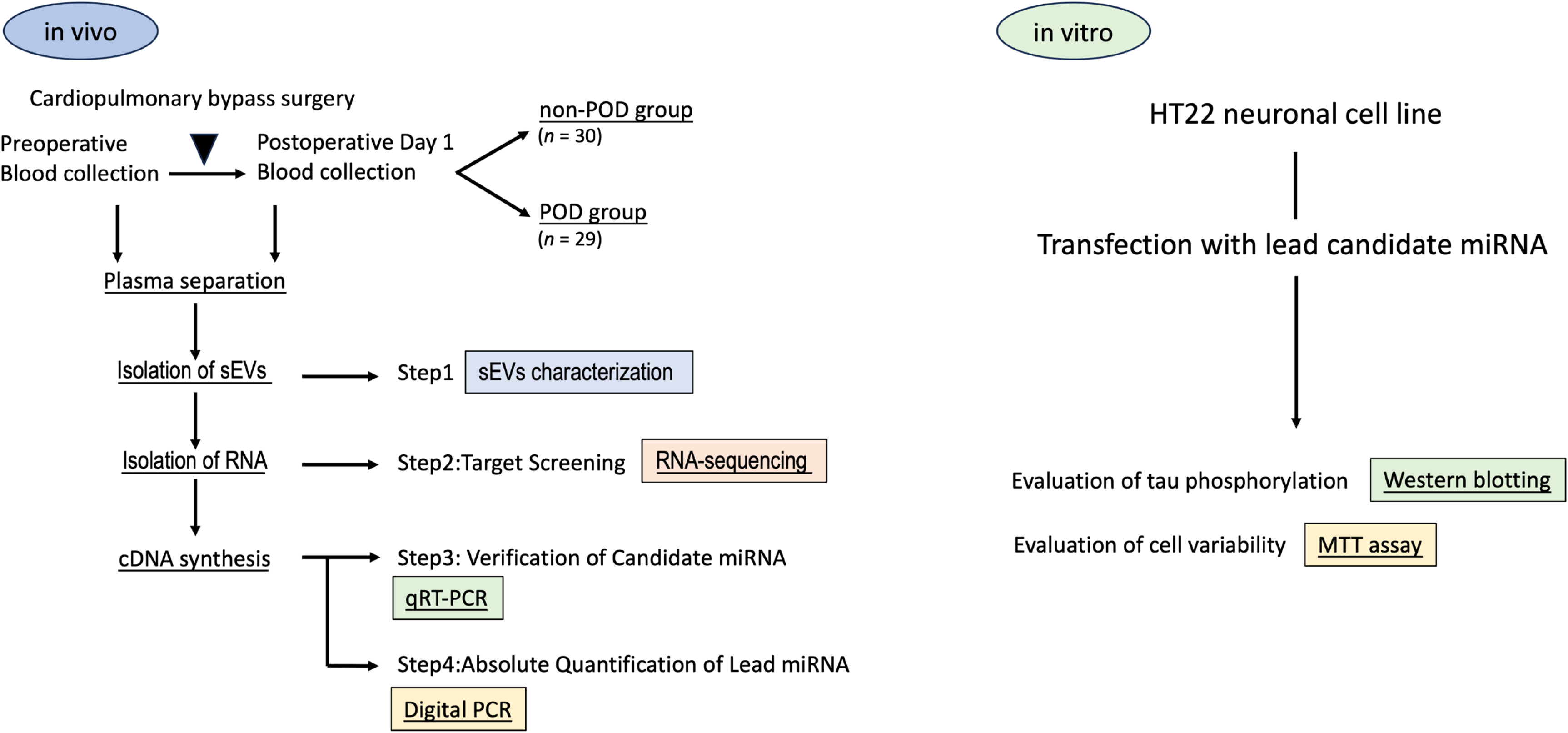
Schematic overview of the study design. The study consisted of an *in vivo* and an *in vitro* arm. The *in vivo* arm involved plasma collection from patients undergoing cardiopulmonary bypass surgery (*n* = 59) at two time points: preoperatively and on postoperative day 1. Samples from patients without postoperative delirium (POD; *n =* 30) and with POD (*n =* 29) were selected in a case-control design from 3 separate cohorts. This was followed by a four-step laboratory workflow: *Step 1*, characterization of isolated small extracellular vesicles (sEVs); *Step 2*, unbiased target screening via RNA-sequencing of sEV-RNA; *Step 3*, validation of candidate microRNAs (miRNAs) using quantitative real-time polymerase chain reaction (qRT-PCR); and *Step 4*, absolute quantification of the lead miRNA using digital PCR (dPCR). The *in vitro* arm investigated the functional consequences of the lead candidate miRNA identified from the *in vivo* analysis. This involved transfection of the HT22 neuronal cell line with the lead candidate miRNA, followed by assessment of tau phosphorylation via Western blotting and cell viability via 3-(4,5-dimethylthiazol-2-yl)-2,5-diphenyltetrazolium bromide (MTT) assay.

### Patient Demographics and Clinical Characteristics

A total of 59 patients undergoing cardiac surgery were included in the final analysis, comprising 30 patients who did not POD (non-POD group) and 29 patients who did (POD group). The demographic and clinical characteristics of the cohort are summarized in **Table 1**. There were no significant differences between the two groups in terms of age, sex, race, or most baseline comorbidities. However, there was a trend toward a higher prevalence of prior stroke (20.7% vs 3.3%, *P* = 0.05) and a longer cardiopulmonary bypass (CPB) time (129 ± 49 min vs 108 ± 51 min, *P* = 0.07) in the POD group.

**Table1.**
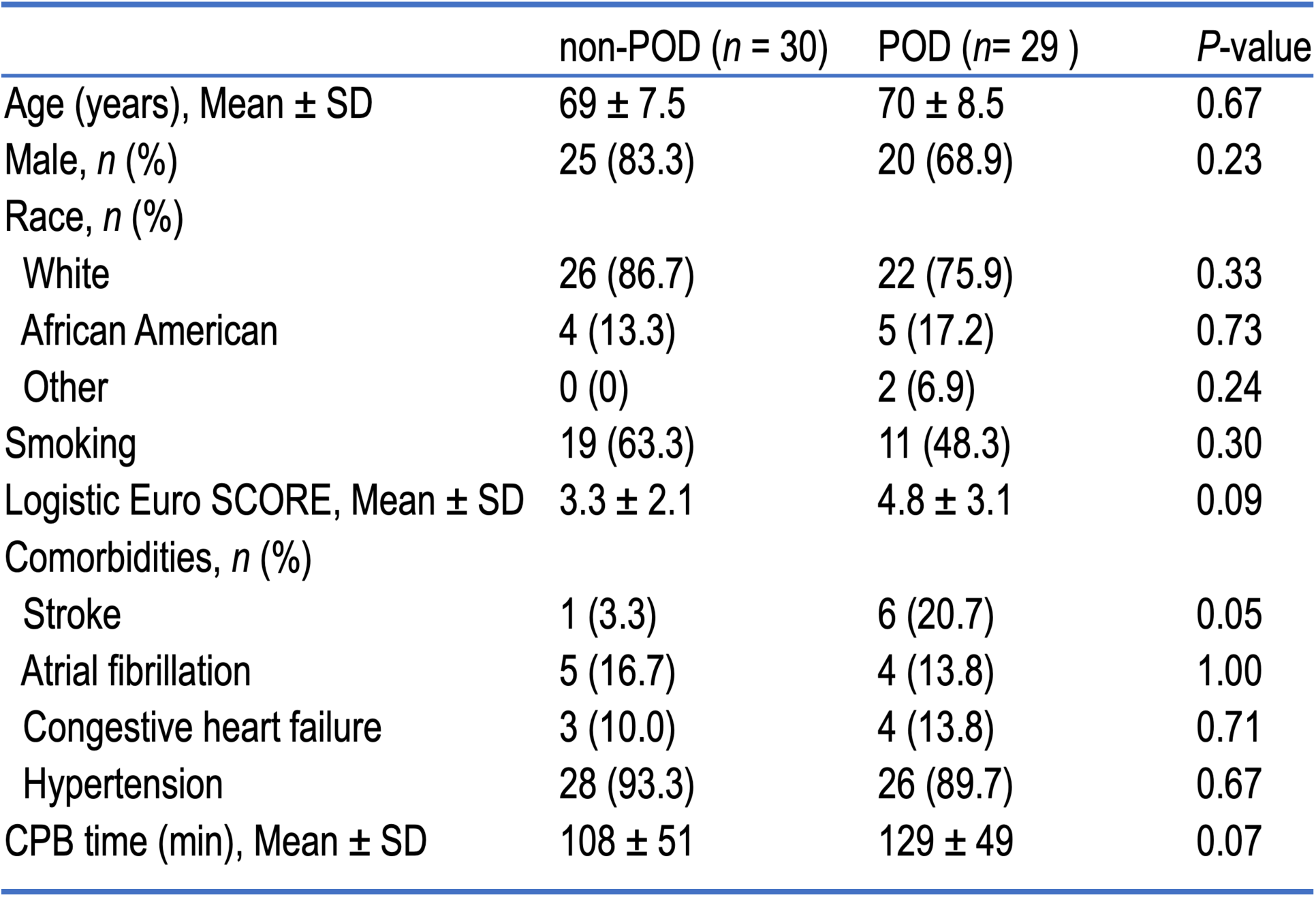
Patient Demographics and Clinical Characteristics Data are presented as mean ± SD for continuous variables and n (%) for categorical variables. p-values for comparisons between non-postoperative delirium (POD) and POD groups were calculated using Mann-Whitney U test for continuous variables and Fisher’s exact test for categorical variables.

|  | non-POD (*n* = 30) | POD (*n*= 29 ) | *P*-value |
| --- | --- | --- | --- |
| Age (years), Mean ± SD | 69 ± 7.5 | 70 ± 8.5 | 0.67 |
| Male, *n* (%) | 25 (83.3) | 20 (68.9) | 0.23 |
| Race, *n* (%) |  |  |  |
| White | 26 (86.7) | 22 (75.9) | 0.33 |
| African American | 4 (13.3) | 5 (17.2) | 0.73 |
| Other | 0 (0) | 2 (6.9) | 0.24 |
| Smoking | 19 (63.3) | 11 (48.3) | 0.30 |
| Logistic Euro SCORE, Mean ± SD | 3.3 ± 2.1 | 4.8 ± 3.1 | 0.09 |
| Comorbidities, *n* (%) |  |  |  |
| Stroke | 1 (3.3) | 6 (20.7) | 0.05 |
| Atrial fibrillation | 5 (16.7) | 4 (13.8) | 1.00 |
| Congestive heart failure | 3 (10.0) | 4 (13.8) | 0.71 |
| Hypertension | 28 (93.3) | 26 (89.7) | 0.67 |
| CPB time (min), Mean ± SD | 108 ± 51 | 129 ± 49 | 0.07 |

### Concentration and Sizes of Isolated sEVs

We first confirmed that sEV number and size were similar between POD groups, which would support the hypothesis that any changes in miRNA levels were due to changes in sEV cargo rather than change in sEV concentration or the size. To establish this, we isolated sEVs from a total of 20 samples (*n =* 5 patients/group, pre- and post-operatively) derived from the dPCR independent validation cohort. The presence of canonical tetraspanin markers (CD63, CD81, and CD9) was confirmed, validating the successful isolation of sEVs (**Figure 2A and 2B**). There were no significant sEV size differences between non-POD group and POD group at both pre-operative and post-operative phase after correction for multiple comparisons (**Figure 2C** and **Supplementary Table 1**). Additionally, there was no significant differences between non-POD and POD groups in terms of CD63, CD81 or CD9 positive sEVs particles concentration (**Figure 2D and Supplementary Table 2**). These results suggest that sEV sizes and concentrations were similar between groups and timepoints, suggesting that any differences in miRNA levels would be due to differential expression within the cargo, rather than changes in the physical characteristics or numbers of the sEVs themselves.

**Figure 2.**
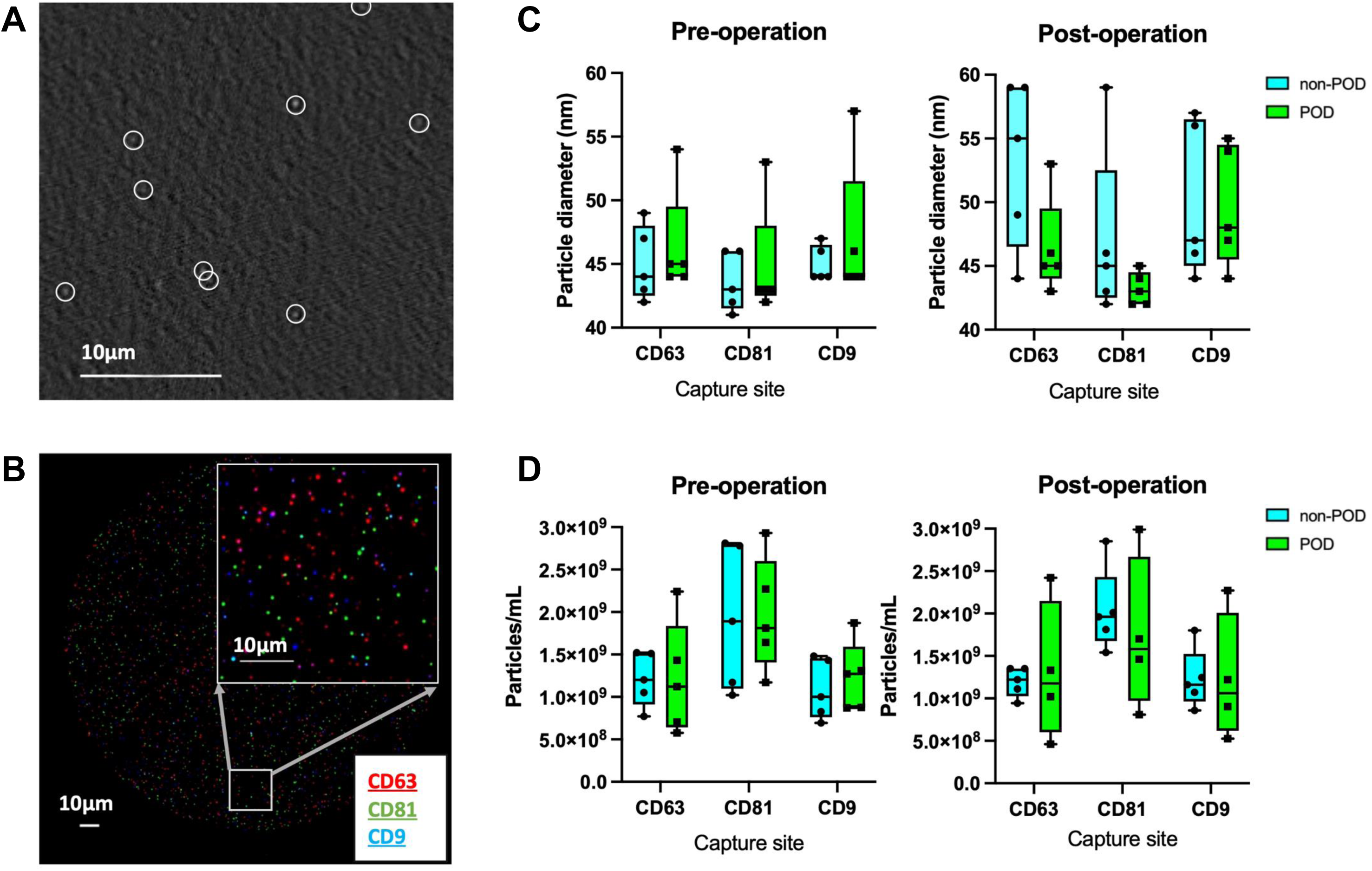
Characterization of sEVs. Immuno-captured sEVs are characterized using (**A**) Single Particle Interferometric Reflectance Imaging Sensor (SP-IRIS) and (**B**) immunofluorescence detection of CD63 (red), CD81 (green) and CD9 (blue), represented by their respective colored puncta. Scale bar, 10 μm. (**C**) Particle size distribution of captured small extracellular vesicles (sEVs) in pre- and post-operative samples. (**D**) Total concentration of captured sEVs on spots targeting CD63, CD81, and CD9. Data in **C** and **D** are presented as box-and-whisker plots showing all individual points (*n* = 5 per group/time point for panel **C**; for panel **D**, *n* = 4-5 per group/time point). The box extends from the 25th to 75th percentiles, the line indicates the median, and the whiskers show the minimum and maximum values. Statistical significance was determined using multiple *t*-tests with False Discovery Rate (FDR) correction.

### Screening and Validation Identifies a Set of Upregulated sEV-miRNAs in POD Patients

To identify potential differences in sEV cargo, we performed an unbiased screen of sEV-miRNA profiles using RNA-sequencing on a discovery cohort of postoperative plasma sEVs (*n =* 6/group; **Figure 3A**). The results revealed 10 miRNAs that were statistically upregulated in the POD group. These target miRNAs were then validated using qRT-PCR in validation cohort (*n =* 10 non-POD, *n =* 8 POD).

**Figure 3.**
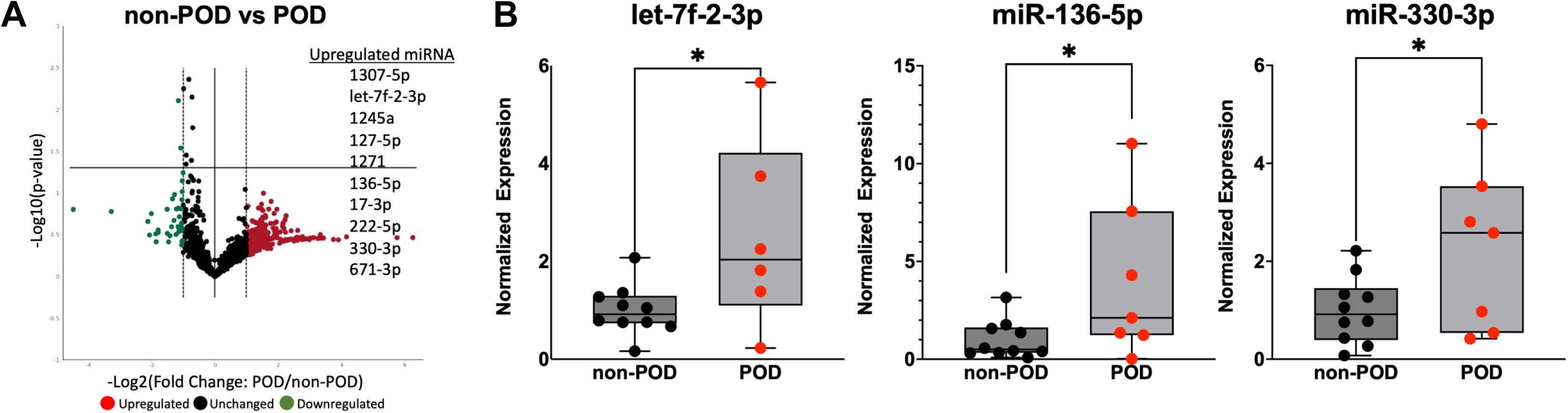
Screening and validation of differentially expressed sEV-miRNAs in postoperative delirium (POD). **(A)** Volcano plot of small RNA-sequencing data comparing small extracellular vesicles (sEV)-miRNA expression between postoperative non-POD (*n =* 6) and POD (*n =* 6) groups. The Y-axis represents statistical significance (-Log_10_ P-value), and the X-axis represents the magnitude of expression change (Log_2_ fold change). Each dot represents a single miRNA. Red dots indicate significantly upregulated miRNAs in the POD group. **(B)** Quantitative real-time polymerase chain reaction (qRT-PCR) validation of the three lead candidate miRNAs in postoperative plasma sEVs. SNO RD61 was used to normalize the miRNA expressions. Data are presented as box-and-whisker plots showing all individual points (*n =* 8-10 for non-POD, *n =* 6-8 for POD). Sample sizes vary between targets due to exclusion of a small number of samples (<5%) due to technical specifications (either undetectable levels or outliers, defined as values exceeding 10-fold of the group mean). The box extends from the 25th to 75th percentiles, the line indicates the median, and the whiskers show the minimum and maximum values. Statistical analysis was performed using Student’s *t*-test. *, *p* < 0.05.

This analysis confirmed that three candidate miRNAs - let-7f-2-3p (*p* = 0.0305), miR-136-5p (*p* = 0.039), and miR-330-3p (*p* = 0.0496) - were significantly upregulated in the POD group (**Figure 3B**). In contrast, the remaining 7 miRNAs did not show significant differences between the two groups (**Supplementary Figure 1**). This multi-step approach thus identified a set of candidate miRNAs whose expression is specifically altered in POD, warranting more precise quantification without any data normalization concerns.

### dPCR Validation confirms upregulation of sEV-miR-330-3p in POD patients

While qRT-PCR is useful for relative validation, absolute quantification is necessary to rigorously evaluate molecular changes. We therefore used dPCR to quantify the exact copy number of the three validated miRNA candidates in a larger cohort (*n =* 20 matched patient pairs/group) (**Figure 4A**). dPCR data revealed that only miR-330-3p showed a significant time-by-group interaction (*p* = 0.0075). Post-hoc tests confirmed that while there was no significant change in the non-POD group, the concentration of miR-330-3p increased significantly from baseline to postoperative day 1 specifically within the POD group (mean increase of 3.80 copies/μL plasma; 95% CI of difference, 0.99 to 6.61; *p* = 0.0087).

**Figure 4.**
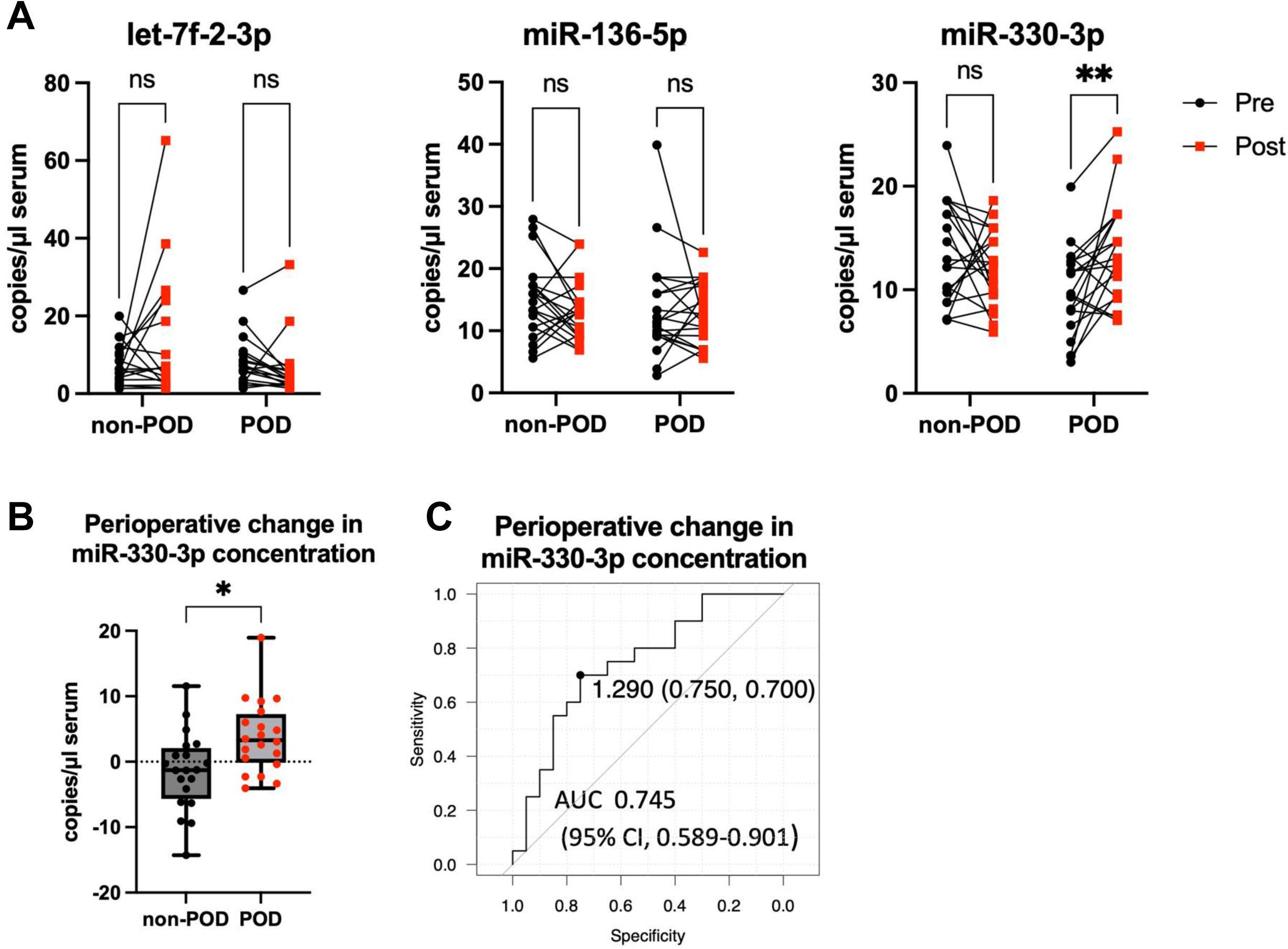
Digital PCR validation of candidate miRNAs and evaluation of miR-330-3p as a predictive biomarker for postoperative delirium (POD). (**A**) Before-after plots show absolute concentrations (copies/μL plasma) of let-7f-2-3p, miR-136-5p, and miR-330-3p at preoperative (Pre) and postoperative (Post) timepoints in non-POD (*n =* 20) and POD (*n =* 20) patients. Each line represents an individual patient. A two-way repeated measures ANOVA revealed a significant time-by-group interaction only for miR-330-3p (*p* = 0.0075). Post-hoc tests confirmed a significant increase in miR-330-3p levels only in the POD group (**, *p* = 0.0087). (**B**) The perioperative change in miR-330-3p concentration (defined as postoperative minus preoperative concentration) was compared between the non-POD and POD groups. Data are presented as a box-and-whisker plot showing all individual points (*n =* 20 per group). The box extends from the 25th to 75th percentiles, the line indicates the median, and the whiskers show the minimum and maximum values. Statistical analysis was performed using a paired *t*-test (*, *p* = 0.0139). (**C**) Receiver operating characteristic (ROC) curve analysis for the ability of the perioperative change in miR-330-3p concentration to predict POD yielded an Area Under the Curve (AUC) of 0.745 (95% Confidence Interval (CI), 0.589 to 0.901). The optimal cut-off value of 1.29 copies/μL plasma provided 75.0% sensitivity and 70.0% specificity.

Based on this specific change we evaluated the perioperative change in miR-330-3p concentration (defined as postoperative minus preoperative concentration) for its ability to distinguish POD from non-POD patients. This change was significantly higher in the POD group compared to the non-POD group (mean difference 5.22 copies/μL plasma; 95% CI, 1.187 to 9.256; *p* = 0.0139; **Figure 4B**). ROC curve analysis yielded an AUC of 0.745 (95% CI, 0.589 to 0.901), indicating a fair diagnostic ability to identify patients who would develop POD. The optimal cut-off value was determined to be 1.29 copies/μL plasma, which provided a sensitivity of 75.0% and a specificity of 70.0% (**Figure 4C**).

### Overexpression of miR-330-3p in Neuronal cells Induces Tau Hyperphosphorylation and Reduces Neuronal Viability

To investigate the functional role of miR-330-3p in neuronal pathology, we overexpressed it in the HT22 neuronal cell line. In accordance with our previous reports ^13, 20^, we confirmed the successful and significant upregulation of miR-330-3p in transfected HT22 cells. Western blot analysis revealed that miR-330-3p overexpression resulted in a marked increase in the phosphorylation of tau protein at both the Ser199 and Ser396 sites, while total tau protein levels remained unchanged (**Figure 5A**). Densitometric quantification confirmed that the ratio of phospho-tau (Ser199) to total tau was significantly elevated in miR-330-3p transfected cells (*p* < 0.0001, **Figure 5B**), as was the ratio of phospho-tau (Ser396) to total tau (*p* < 0.001, **Figure 5C**). No significant differences in total tau expression were observed among the groups (**Figure 5D**).

**Figure 5.**
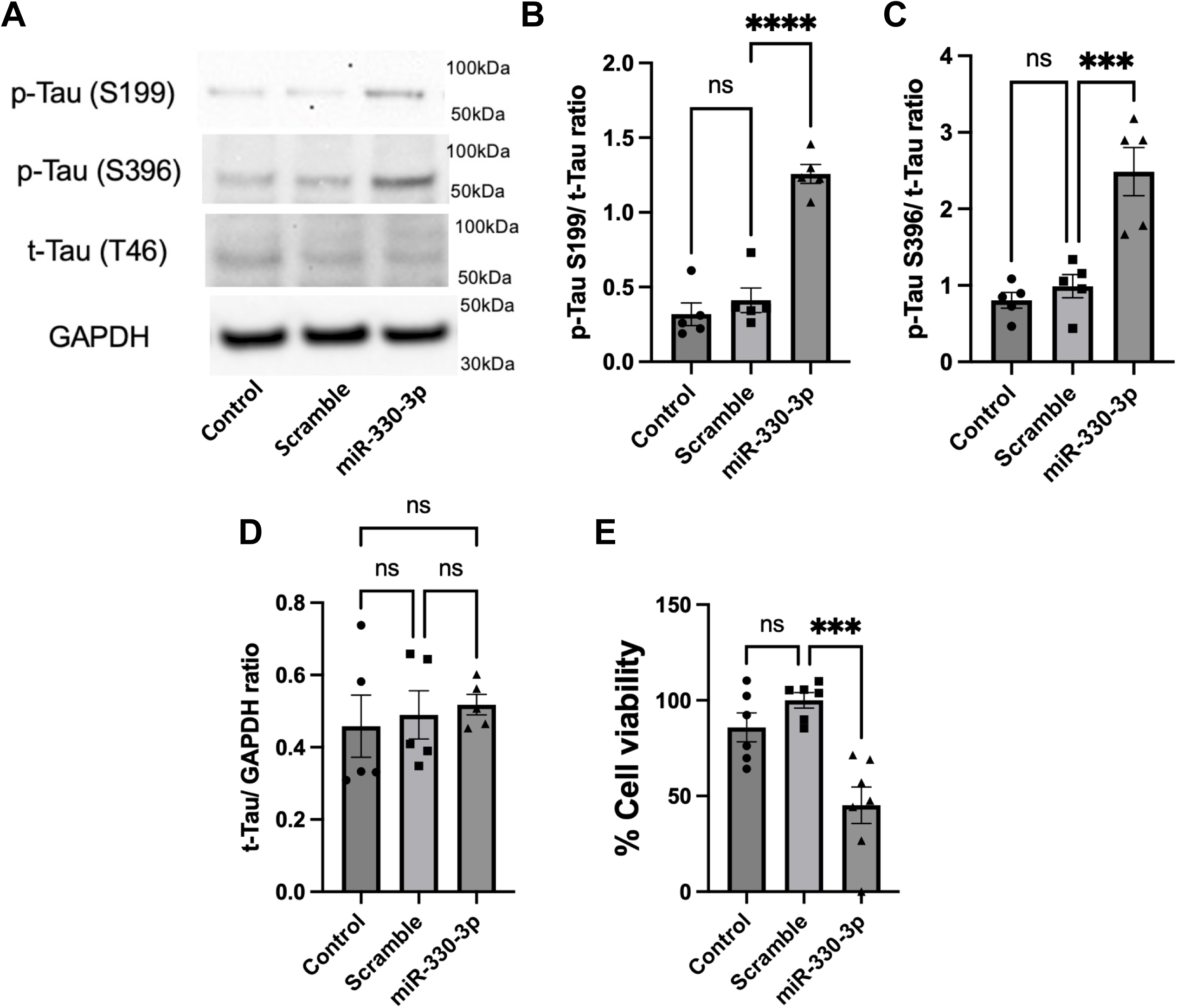
Overexpression of miR-330-3p induces tau hyperphosphorylation and reduces viability in HT22 neuronal cells. **(A)** Representative western blot images showing expression of phospho-tau (p-Tau) at Ser199 and Ser396, total tau (t-Tau), and GAPDH (loading control) in the three experimental groups. **(B-D)** Densitometric quantification of Western blots for the ratio of p-Tau S199 to t-Tau (**B**), p-Tau S396 to t-Tau (**C**), and t-Tau to Glyceraldehyde-3-phosphate dehydrogenase (GAPDH; **D**) (*n* = 5 / group). **(E)** 3-(4,5-dimethylthiazol-2-yl)-2,5-diphenyltetrazolium bromide (MTT) assay shows a significant reduction in the percentage of viable cells in the miR-330-3p mimic-transfected group (*n* = 6 / group). Data in bar charts are presented as mean ± SEM with all individual data points shown from at least three independent experiments. Statistical analysis was performed using one-way ANOVA with Tukey’s post-hoc test. \*\*\**P* < 0.001, \*\*\*\**P* < 0.0001; ns, not significant.

Finally, to assess the cytotoxic effects of miR-330-3p, we performed an MTT assay. The results demonstrated that miR-330-3p overexpression significantly decreased neuronal viability compared to control and scramble groups (*P* < 0.001, **Figure 5E**), indicating that elevated levels of miR-330-3p are detrimental to neuronal cell survival. These findings provide direct *in vitro* evidence for a potential pathogenic role of miR-330-3p, connecting the molecule identified in our clinical samples to a key pathway of neuronal injury.

## Discussion

In this study, we combined a prospective clinical investigation with *in vitro* mechanistic approach to explore the role of circulating sEV-cargo miRNAs in the pathogenesis of POD. Our principal finding is that a perioperative increase in sEV-cargo miR-330-3p is associated with patients who develop POD. Furthermore, we provide novel evidence that miR-330-3p can directly induce tau hyperphosphorylation and reduce neuronal viability, suggesting a potential molecular pathway linking the systemic surgical response to CNS pathology. This work also introduces a robust methodology for identifying and functionally validating sEV-miRNA signals in perioperative neurocognitive disorders.

A strength of our approach is the focus on sEV-cargo miRNAs. Circulating miRNA studies have historically been challenged by variability in methods and an underappreciation of the miRNA’s cellular source, often reflecting contamination from blood cells rather than a true pathological signal.^21, 22^ By focusing on sEVs, which protect their cargo from degradation and provide a more enriched source of biomarkers compared to whole plasma, our methodology provides a more robust and reproducible strategy.^23^ This approach is particularly novel in the context of perioperative neurocognitive disorders, where the specific molecular messengers linking the peripheral surgical site to the brain have remained elusive. Since miR-330-3p is highly expressed in neurons and systemic inflammation and cytokine release has been reported to activate neuronal miR-330-3p expression^24^, a plausible mechanism is that peripheral surgical trauma induces neuroinflammation, which in turn upregulates neuronal miR-330-3p expression and promotes its incorporation into sEVs. It has been shown that miR-330-3p can directly bind to the 3’-UTR of *ULK1* (Unc-51-like kinase 1) mRNA, a crucial mediator in neuronal survival and neurotoxicity. ^25^

We specifically examined perioperative changes in miRNA, since the absolute concentration and cargo of circulating sEVs are known to be highly variable between individuals, influenced by a multitude of physiological and pre-analytical factors.^26^ This inherent variability makes the establishment of a universal reference range exceedingly difficult. By using each patient as their own baseline and focusing on the dynamic change in response to surgery, our study mitigates these confounding factors to isolate a disease-specific signal. The ROC analysis for this marker yielded an AUC of 0.745 for prediction of POD. While this indicates a modest predictive ability for a standalone test, future studies will need to determine whether multivariable risk prediction models that incorporate a number of biologic signals (including other miRNA) can add a dynamic dimension to static preoperative clinical risk factors.^5^

The most significant contribution of this study is the elucidation of a potential pathogenic mechanism. Our *in vitro* data are the first to demonstrate that miR-330-3p can directly induce hyperphosphorylation of tau at Ser199 and Ser396 (**Figure 5B and 5C**). This is a crucial finding, as abnormal tau phosphorylation is associated with cognitive impairment through synaptic dysfunction and neuronal loss ^27^, and is a hallmark of neurodegenerative diseases.^28^ An increasing body of evidence directly implicates tau pathology in the acute neuronal injury associated with delirium.^29^ Our findings add to a growing body of literature on the role of miR-330-3p in neurological dysfunction. While miR-330-3p has been extensively studied for its role in cancer, including glioblastoma and brain metastasis of non-CNS cancer, its direct involvement in neurocognitive disorders like Alzheimer’s Disease and Parkinson’s Disease has also been documented.^30–33^ Importantly, miR-330-3p has been previously reported as a potential biomarker for age-associated mild cognitive impairment.^34^ Our findings on the neurotoxic role of miR-330-3p are further supported by recent preclinical studies. For example, Xu and colleagues demonstrated that an anesthetic agent induced neurotoxicity in rats by upregulating miR-330-3p in the hippocampus, and that inhibiting miR-330-3p was neuroprotective.^35^

The concept of periphery-to-brain communication via sEVs provides a compelling biological framework for our findings. Macrophage-derived sEVs can transport inflammatory cargo that impacts CNS function.^36^ We demonstrated that the physical properties of the sEVs themselves did not differ between the groups, strongly suggesting that the critical difference lies within their molecular cargo. The systemic inflammation from cardiac surgery may trigger the release of neuronal sEVs with altered miRNA content from peripheral tissues ^37^, which then cross a compromised blood-brain barrier to exert their effects.^38^ This aligns with the MISEV2023 guidelines, which emphasize the importance of characterizing both vesicles and their cargo.^39^

This study has several strengths, including using samples from multiple cohorts with rigorous delirium assessments, having derivation and validation approaches, using assays that provide absolute concentrations, and translating findings from human cohorts into *in vitro* assays. However, we acknowledge limitations. The sample size necessitates validation in larger, multi-center cohorts. The cellular origin of the sEVs remains unknown, and our *in vitro* findings do not definitively prove causality *in vivo*. Future research should focus on validating miR-330-3p in diverse surgical populations and incorporating it into multivariable prediction models. Mechanistically, identifying the direct mRNA targets of miR-330-3p that regulate tau phosphorylation is a critical next step. Animal models will also be essential to establish causality and to explore the exciting therapeutic possibility suggested by our findings: that inhibiting this pathway with a specific miR-330-3p antagonist could represent a novel intervention to prevent POD.

In conclusion, this study identifies an increase in circulating sEV-cargo miR-330-3p in patients with POD. We provide a novel mechanistic link to tau pathology, suggesting that sEV-mediated miRNA transfer may participate in the development of perioperative neurocognitive disorders. This work not only provides a promising biomarker candidate to incorporate into risk stratification models but also uncovers a new potential therapeutic target to protect brain health after surgery.

## Author Contributions

TF, SC, AM and HT performed the experiments, analyzed the data, helped to draft the article and revise it critically for important intellectual content, provided final approval of the version to be published; and has agreed to be accountable for all aspects of the work.

AY made a substantial contribution to conception and design of the article, revise it critically for important intellectual content, provided final approval of the version to be published; and has agreed to be accountable for all aspects of the work.

CH obtained informed consent and collected patient samples for the study, contributed to drafting the manuscript and critically revised it for important intellectual content, and provided final approval of the version to be published.

CHB and SD made a substantial contribution to conception and design of the experiments, reviewing the data and helped with the analysis, helped to draft the article and revise it critically for important intellectual content, provided final approval of the version to be published; and has agreed to be accountable for all aspects of the work.

## Acknowledgements

None

## Declaration of Interests

None.

## Funding

SD acknowledges grants from the U54AG062333 and U18TR003780 from National Institutes of Health and by American Heart Association grants TPA 970850 and 23DIVSUP1057308. CB has funding from NIH R01HL092259.

## Nonstandard Abbreviations and Acronyms

miR or miRNA: microRNA
EV: Extracellular Vesicles
sEV: small Extracellular Vesicles
qRT-PCR: Quantitative real-time polymerase chain reaction
dPCR: Digital PCR
RNA-Seq: RNA Sequencing
CI: Confidence Interval
POD: Postoperative Delirium
CPB: Cardiopulmonary bypass
MTT 3-(4,5-dimethylthiazol-2-yl)-2: 5-diphenyltetrazolium bromide
Tau at Ser396: Tau protein at Serine 396 site
Tau at Ser199: Tau protein at Serine 199 site

## Notes

### Competing Interest Statement

The authors have declared no competing interest.

